# Reference genome sequence of the solitary bee *Camptopoeum friesei* Mocsáry, 1894 (Hymenoptera, Andrenidae)

**DOI:** 10.1101/2023.08.27.555015

**Authors:** Eckart Stolle, Nadège Guiglielmoni, Joseph Kirangwa, Sandra Kukowka, Tobias Meitzel, Ann M. Mc Cartney, Stefanie Heilmann-Heimbach, Kerstin Becker, Karl Köhrer, Astrid Böhne

## Abstract

Bees are major pollinators of flowering plants and thus are important ecosystem service providers for natural habitats and crops. Evolution led to a wide range of adaptations in behaviors, morphology and ecological traits. Many plants rely on specialized bee species for pollination events, and so this interdependence can make them increasingly vulnerable to ongoing threats of habitat loss and pesticide exposure. Studying the genomes of bee species across different life histories and ecological specializations can help understand the evolution of these traits more generally, but also inform conservation efforts for *Camptopoeum friesei* specifically.

Here, we present the reference genome of the solitary bee *Camptopoeum friesei* (Arthropoda; Insecta; Hymenoptera; Andrenidae).

*C. friesei* is highly dependent on steppe habitats where it nests in saline soils. Further, it is highly specialized (oligolectic) on a few Asteraceae: *Centaurea* and *Cirsium*, in particular on *Centaurea stoebe*. As a consequence of its high specialization level, it is of its ecological niche with an extremely scattered and rare habitat, *C. friesei* is highly threatened in central Europe, albeit local aggregations can be rich in individuals.

The high-quality genome assembly for the colourful bee *Camptopoeum friesei* was generated using long-read PacBio HiFi in combination with chromatin conformation capture (Hi-C) sequencing. The genome spans 367.7 megabases (Mb), N50 of 25.2 Mb. The majority of the assembly is scaffolded into 10 chromosomes and harbours ∼40% repeats.

**Species taxonomy:** Eukaryota; Opisthokonta; Metazoa; Eumetazoa; Bilateria; Protostomia; Ecdysozoa; Panarthropoda; Arthropoda; Mandibulata; Pancrustacea; Hexapoda; Insecta; Dicondylia; Pterygota; Neoptera; Endopterygota; Hymenoptera; Apocrita; Aculeata; Apoidea; Anthophila; Andrenidae; Panurginae; Panurgini; *Camptopoeum friesei* Mocsáry, 1894 (NCBI:txid2918745)

## Introduction

Bees are major ecosystem service providers in both natural and anthropogenic habitats. Akin to most other important pollinators bees play a vital role in ecosystem functioning and are, due to their interdependence with plants and nesting habitats, especially threatened by habitat loss, pesticide exposure and climate change (Klein et al. 2007; Steffan-Dewenter et al. 2005). Consequently, declines in bee populations threatens the integrity of ecosystem function and food security.

The vast majority of bees are solitary while different levels of sociality evolved in multiple subclades, most notably the honeybees, bumblebees and stingless bees. With almost 20,000 described species worldwide (Michener 2000; Danforth et al. 2019), bees are successful, speciose and essential components of the terrestrial ecosystem due to their important role as pollinators (Klein et al. 2007; Allen-Wardell et al. 1998). Larvae of almost all bee species ingest a diet of pollen combined with nectar or floral oils.

Given their importance for commercial and ecological pollination and their social lifestyle, honeybees and bumblebees are among the best studied and accessible insects. The genome of the honeybee was among the first genomes available for any insect (Honeybee Genome Sequencing Consortium 2006). Since then, most of the genomes of bees have been generated with a focus on the evolution of sociality (e.g.,(Elsik et al. 2014; Sadd et al. 2015; Jones et al. 2023; Kapheim et al. 2015; Kocher et al. 2013; Rehan et al. 2016; Crowley et al. 2023)). Only recently additional solitary species became available (e.g. (Ferrari et al. 2023; Zhou et al. 2020; Zhang et al. 2022; Falk, Monks, et al. 2022; Falk, University of Oxford and Wytham Woods Genome Acquisition Lab, et al. 2022; Boyes et al. 2023)).

However, it is critical to broaden sequencing efforts to include species from under-represented bee taxa. Expanding genomic exploration beyond social bee species can facilitate a greater understanding of the diverse life histories and ecological adaptations of all bee species. Broader comparisons can not only help to understand the evolution of sociality from solitary ancestors (Robinson et al. 2005; Toth & Robinson 2007), but also can shed light on the evolution of ecological adaptations such as habitat niche, flower specialization, distribution and climate adaptation. Furthermore, many understudied bee taxa contribute important pollination services. Many bee species have a high degree of specialization to specific plant families or even species, hence can be under more pressure from climate change and habitat loss. While generalist bee species often have more opportunities to adapt, specialists are overrepresented among highly threatened species. Specialists hence suffer often steeper population declines and require higher, or more specific conservation needs. As a result, examining a diverse set of bee genomes and biologies is critical for conservation efforts and tackling the issues they may face in the face of environmental pressures (De Palma et al. 2015).

The solitary bee *Camptopoeum friesei* is native to Europe and belongs to the Andrenidae bee family. It is highly dependent on steppe-like habitats with saline clay soils where it nests underground and it is highly specialized (oligolectic) on *Centaurea* and *Cirsium* plants (Asteraceae) (Jansen & Saure 2021; Koppitz et al. 2017). In central Europe it occurs only in secondary (salt mining silt deposit) habitats in central Germany and in lower Austria, i.e. albeit local aggregations can be large, the populations are highly isolated and comprise only of a very small area (Koppitz et al. 2017; Jansen & Saure 2021; Dorn 1969; Stolle 2014). As a result of this, it is considered highly threatened in Germany (Westrich et al. 2011; Saure & Stolle 2016; Saure 2020). The only population in central Germany was first mentioned 1901 by H. Friese (Friese 1901) based on a communication with Alfken who received a male collected near Eisleben on 23. August 1891 (Alfken 1912). However, since *C. friesei* was only described 1894 (Mocsáry 1894), the individual from 1891 was identified as *C. frontale* which was henceforth used as species identity of this population (Dorn 1969; Koppitz et al. 2017; Stolle 2014) until clarification in 2021 (Jansen & Saure 2021).

The reference genome of *Camptopoeum friesei* generated herein, through a combination of long-read PacBio HiFi sequencing and chromatin conformation capture (Hi-C), will facilitate comparative genomic approaches for the identification of genetic signatures and molecular mechanisms that underlie ecological specialization and will facilitate understanding long-term consequences of isolation in an insect population. The reference genome provides the basis of population-wide insights into genetic diversity, genetic load, demography and population connectivity, hence establishing genomic information to assist conservation efforts.

## Materials and Methods

### Sample acquisition and DNA extraction

Haploid male bees were gathered from a population near Bernburg (Germany) and easily classified to the species level by eye. A single haploid male bee was selected, and high-molecular-weight (HMW) DNA was extracted from it using the NEB Monarch HMW DNA Extraction Kit by New England Biolabs. Quality control (QC) of input DNA for sequencing is important for assessing the integrity and average size of a sample and can help determine the cutoff size for size selection. The quality of the DNA was assessed using the Nanodrop 2000 spectrometer and the Qubit v4 fluorometer with the dsDNA HS assay kit from Invitrogen, Carlsbad, CA, USA. Additionally, the integrity and size of the DNA fragments were confirmed using the Agilent Femto Pulse System (automated pulsed-field capillary electrophoresis system for sizing high molecular weight (HMW) genomic DNA (gDNA). The extracted high-molecular-weight DNA was stored at +4 °C prior to HiFi library preparation and long read sequencing.

### RNA extraction

RNA was extracted from two separate individuals (male and female) with the Qiagen RNA Micro and Mini kit according to manufacturer’s instructions. RNA quality and quantity was determined using the Nanodrop 2000 and stored at -80°C.

### Long read sequencing

HiFi libraries were prepared with the Express 2.0 Template kit (Pacific Biosciences, Menlo Park, CA, USA) and sequenced on a Sequel II/Sequel IIe instrument with 30h movie time. HiFi reads were generated using SMRT Link (v10; (Pacific Biosciences, Menlo Park, CA, USA) with default parameters.

### Hi-C sequencing

A half individual was cross-linked in 3% formaldehyde for 1 hour at room temperature. The reaction was quenched with glycine at a final concentration of 250 mM. The individual was flash-frozen in liquid nitrogen and ground using a pestle. Hi-C libraries were prepared using the Arima-HiC+ kit (Arima Genomics, Carlsbad, CA, USA) according to manufacturer’s instructions, and subsequently paired-end (2×150 bp) sequenced on a NovaSeq 6000 instrument (Illumina, San Diego, CA, USA).

### RNA sequencing

Library preparation for full-length mRNASeq was performed using the NEB Ultra II Directional RNA Library Prep with NEBNext Poly(A) mRNA Magnetic Isolation Module and 500 ng total RNA as starting material. Sequencing was performed on an Illumina NovaSeq 6000 device with 2×150bp paired-end sequencing protocol and >50M reads per sample.

### Genome assembly

*k*-mers in the PacBio HiFi reads were analyzed using jellyfish v2.3.0 and GenomeScope2 (Ranallo-Benavidez et al. 2020) with parameters *k*=27 and l=30. PacBio HiFi reads were assembled using hifiasm v0.16.1-r375 (Cheng et al. 2021) with parameter -l 2. Hi-C reads were trimmed using cutadapt v1.15, mapped using bowtie2 2.5.0 (Langmead & Salzberg 2012) and processed using hicstuff v3.1.5 (Matthey-Doret et al.) with parameters -e DpnII,HinfI -m iterative. Small contigs with low to no Hi-C contacts were removed. The aligned reads were further processed using the Arima mapping pipeline with the script two_read_bam_combiner.pl (https://github.com/ArimaGenomics/mapping_pipeline) and Picard v2.8.1 (http://broadinstitute.github.io/picard/) modules AddOrReplaceReadGroups and MarkDuplicates with parameters ASSUME_SORTED=TRUE VALIDATION_STRINGENCY=LENIENT REMOVE_DUPLICATES=TRUE. The output was provided to PretextMap v0.1.9 (https://github.com/wtsi-hpag/PretextMap) and the contact map was used with PretextView v0.2.5 (https://github.com/wtsi-hpag/PretextView) to manually reconnect two contigs with strong contacts using the GRIT Rapid Curation workflow (https://gitlab.com/wtsi-grit/rapid-curation). Ortholog completeness was assessed using the Benchmarking Universal Single-Copy Orthologs (BUSCO) tool v.5.0.0 against the Hymenoptera odb10 lineage (5,991 orthologs) (Waterhouse et al. 2018). PacBio HiFi reads were mapped to the final scaffolds using minimap2 v2.24-r1122 (Li 2018) with parameter -ax map-hifi and the mapped reads were sorted with SAMtools v1.11 (Li et al. 2009). The scaffolds were aligned against the nucleotide database using the Basic Local Alignment Search Tool (BLAST) v2.6.0 (Altschul et al. 1990) with parameters -outfmt “6 qseqid staxids bitscore std sscinames scomnames” -max_hsps 1 -evalue 1e-25. The outputs of minimap2, BLAST, and BUSCO were provided as input to Blobtools2 (Challis et al. 2020) and contigs flagged as proteobacteria were subsequently removed. *k*-mer completeness was evaluated using KAT v2.4.2 (Mapleson et al. 2017) and the module kat comp with default parameters. BUSCO and hicstuff were run on the final assembly as previously described and the Hi-C contact map was visualized using hicstuff view with parameter -b 2000. Repeats were identified de novo using EDTA (Ou et al. 2019).

## Results

The genome of *Camptopoeum friesei* (Figure 1) was assembled using 32.3 Gigabases (Gb) of PacBio HiFi reads with an N50 of 13.1 kilobases (kb) and 204 millions pairs of 150-basepairs (bp) Hi-C reads. The draft assembly was generated using hifiasm; these contigs included proteobacteria contaminants (Figure 2) and contigs with low Hi-C coverage, which were subsequently removed. After curation, the final assembly has a size of 367.7 Megabases (Mb) and an N50 of 25.2 Mb (Table 1). 12 scaffolds have a size ranging from 11.0 to 37.8 Mb and 10 of these can be identified in the Hi-C contact map (Figure 3). The BUSCO score (against Hymenoptera odb10) reaches 97.3% (including 0.2% duplicated orthologues), the QV score is estimated to 69.1 and the *k*-mer plot shows no artefactual duplications (Figure 4). All these evaluations support the high quality of this assembly despite unusual Hi-C patterns for two scaffolds, which require further investigation.

**Table 1.** Overview for the reference genome sequence assembly of *Camptopoeum friesei*.

|  | **bp** | **n** |
| --- | --- | --- |
| total length / scaffolds | 367718021 | 48 |
| average length | 7660792.1 |  |
| largest scaffold | 37766881 |  |
| n50 | 25206928 | 6 |
| n60 | 24095866 | 8 |
| n70 | 22801990 | 9 |
| n80 | 13191795 | 11 |
| n90 | 2767723 | 19 |
| n100 | 342844 | 48 |
| N_count / gaps | 200 | 1 |

**Figure 1.**
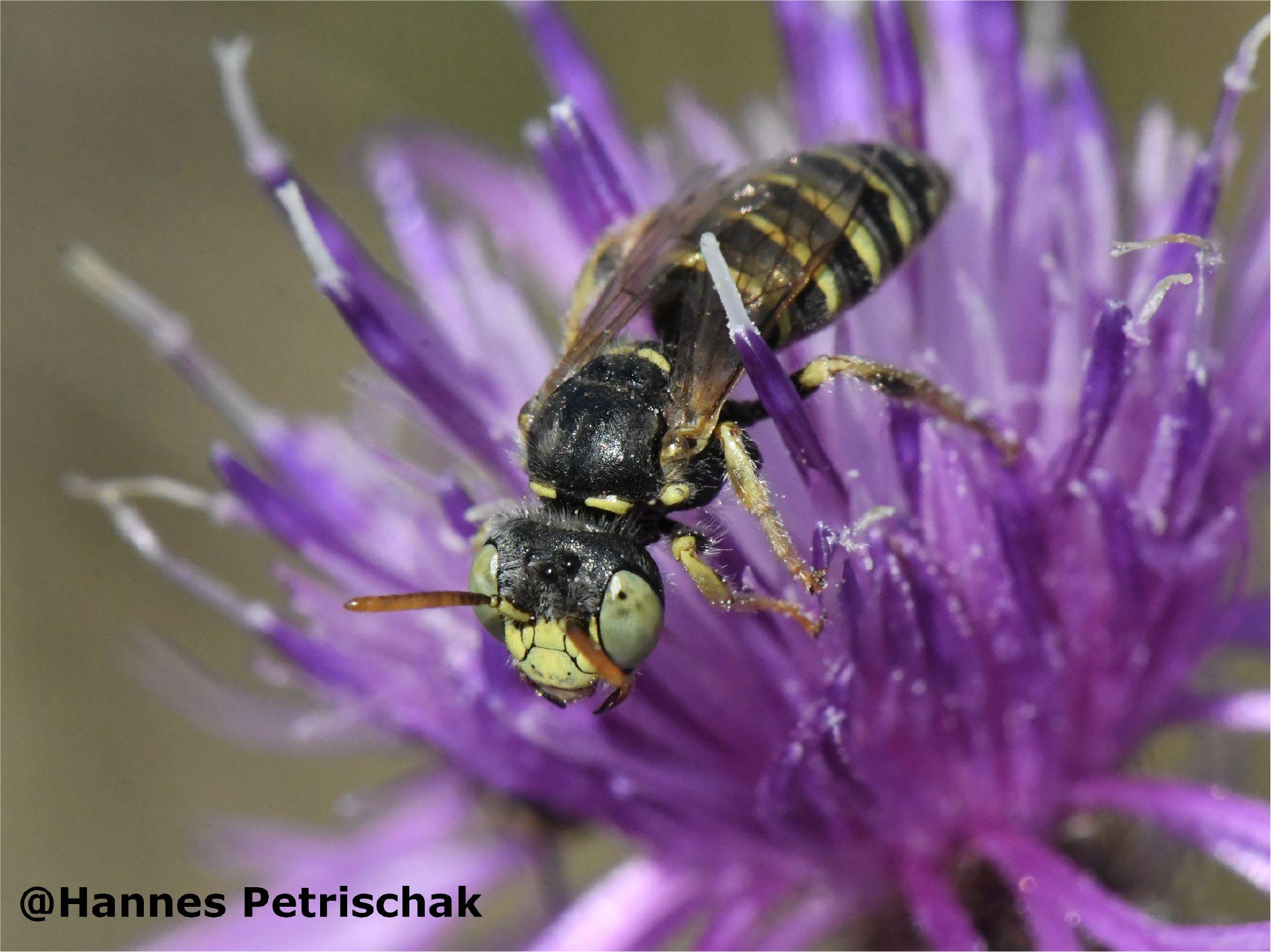
Male of the steppe mining bee *Camptopoeum friesei*. Phooto: Hannes Petrischak

**Figure 2.**
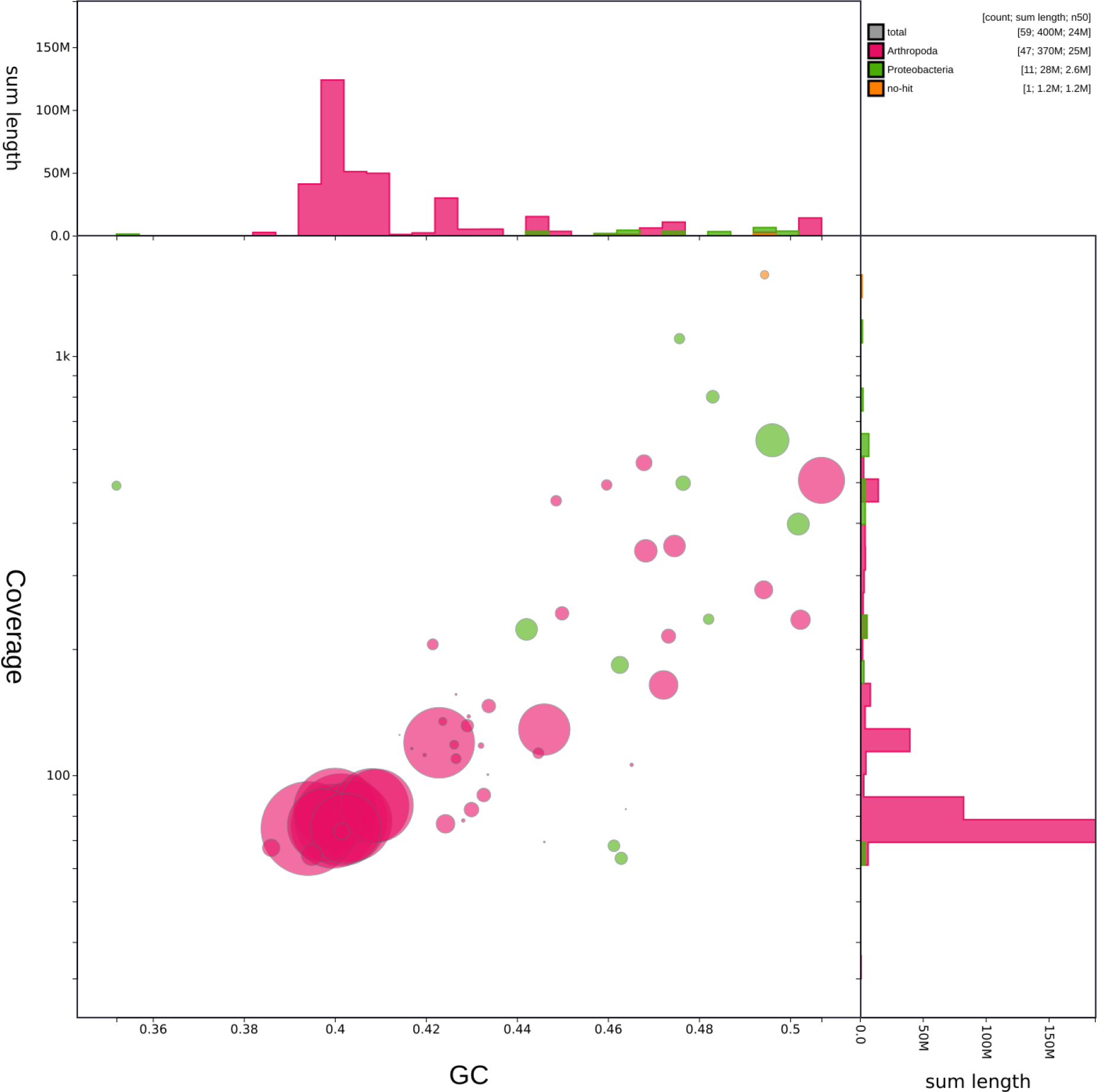
BlobTools analysis. Contaminant contigs (flagged as Proteobacteria) were removed from the final assembly.

**Figure 3.**
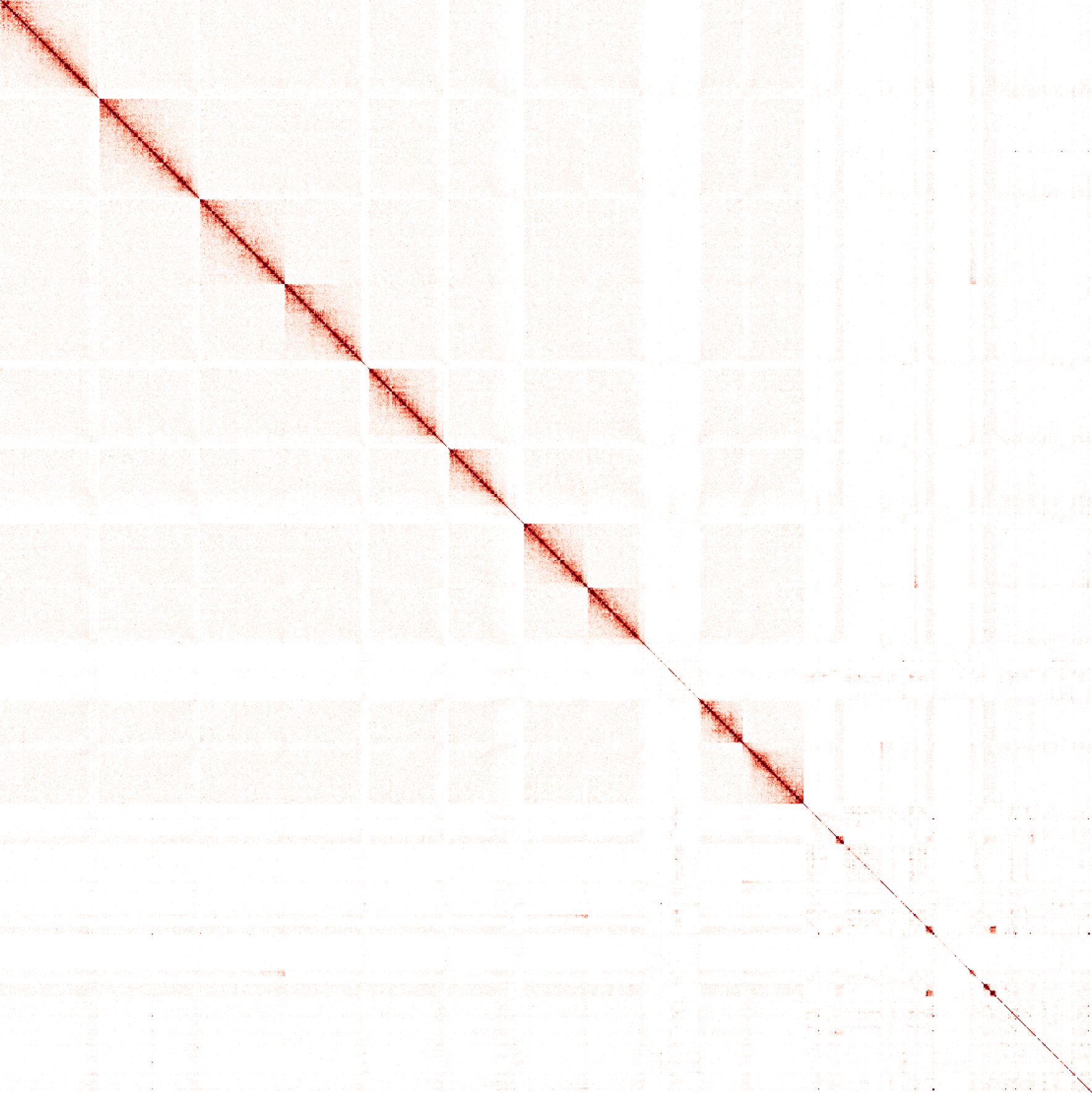
Hi-C contact map of the final assembly.

**Figure 4.**
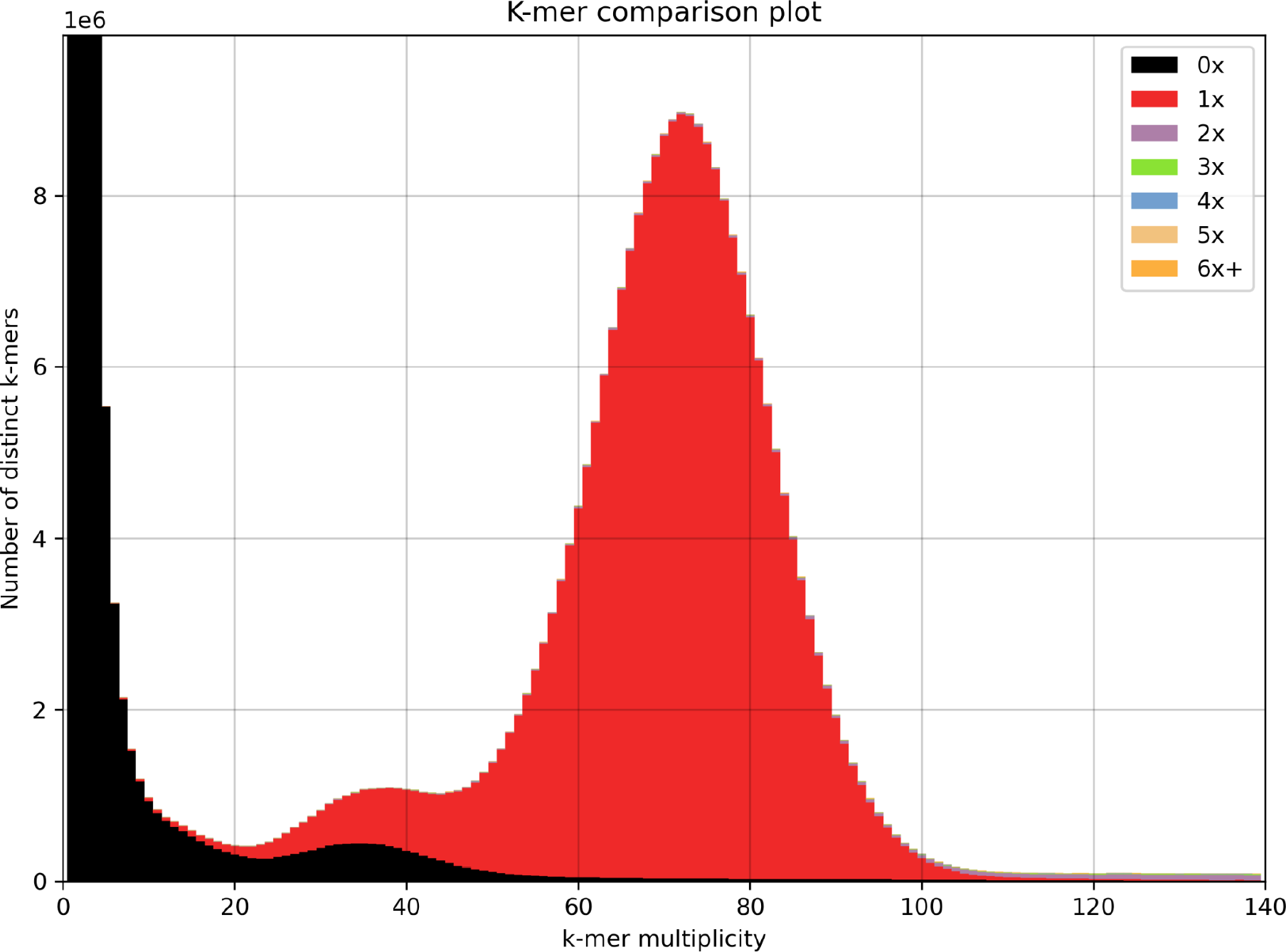
*k*-mer comparison of the PacBio HiFi reads v. the final assembly.

De novo repeat annotation identified 40.05% as repetitive, with 11.34% Terminal inverted repeat DNA transposons, 9.24% LTR retrotransposons and 4.03% Helitrons being the most common interspersed repeats (Table 2).

**Table 2.** Repeats in the reference genome sequence assembly of *Camptopoeum friesei*.

| **Class** |  | **count** | **length (bp)** | **proportion of the genome** |
| --- | --- | --- | --- | --- |
| LINE | unknown | 2278 | 2311107 | 0.63% |
| LTR | Copie | 6597 | 4517222 | 1.23% |
| LTR | Gypsy | 31744 | 27400741 | 7.45% |
| LTR | unknown | 5539 | 2075028 | 0.56% |
| TIR | CACTA | 21635 | 4874817 | 1.33% |
| TIR | Merlin | 24 | 6239 | 0.00% |
| TIR | Mutator | 49485 | 13403668 | 3.65% |
| TIR | PIF_Harbinger | 9202 | 2342761 | 0.64% |
| TIR | Tc1_Mariner | 3746 | 915736 | 0.25% |
| TIR | hAT | 50507 | 18970518 | 5.16% |
| TIR | piggyBac | 203 | 161543 | 0.04% |
| TIR | polinton | 719 | 976168 | 0.27% |
| nonLTR | DIRS_YR | 139 | 128963 | 0.04% |
| nonLTR | helitron | 61181 | 14812347 | 4.03% |
| repeat_region |  | 315671 | 54367798 | 14.79% |
| **total**<br>**interspersed** |  | **558670** | **147264656** | **40.05%** |

## Data availability

Submission to ENA in progress, accession numbers pending.

## Acknowledgments

This genome project is part of the DeRGA pilot study (https://www.erga-biodiversity.eu/pilot-253 project) and we acknowledge coordination and support by Dr. Philipp Schiffer and Prof. Dr. Ann-Marie Waldvogel (University of Cologne). Development of the pilot ERGA Data Portal (https://portal.erga-255 biodiversity.eu) was funded by the European Molecular Biology Laboratory. We thank the West German Genome Center (WGGC) and the Cologne Center for Genomics (CCG) for library generation and quality control as well as their support for PacBio Hifi, Hi-C and RNA sequencing. We thank Dr. H. Petrischak (Heinz Sielmann Stiftung, Wustermark) for providing pictures of *C. friesei*.

